# Biogeographic affinities, phylogenetic relationships, and diversity of ant species of the genus *Camponotus* (Hymenoptera: Formicidae) present in Chile

**DOI:** 10.1101/2025.11.30.691467

**Authors:** Benjamín Arenas-Gutierrez, Carlos P. Muñoz-Ramírez

**Affiliations:** Programa de Magíster en Ciencias mención Entomología, Instituto de Entomología, Universidad Metropo-litana de Ciencias de la Educación, Av. José Pedro Alessandri 774, Ñuñoa, Santiago, Chile; Vicerrectoría de Investigación e Innovación, Universidad Arturo Prat. Dirección. Avenida Arturo Prat 2120, Iquique, Chile; Instituto de Entomología, Universidad Metropolitana de Ciencias de la Educación, Av. José Pedro Alessandri 774, Ñuñoa, Santiago, Chile

**Keywords:** Ants, Biogeography, Camponotini, COI, Cryptic Species, Evolution, Formicidae, Phylogeny

## Abstract

The biogeographic relationship of the entomofauna present in Chile has been studied in different groups (e.g. Diptera, Coleoptera, Plecoptera), finding in many cases a possible Gondwanic origin, with a greater relatedness with the Austral/Oceanic entomofauna. In the case of ants, specifically *Camponotus* (Hymenoptera, Formicidae) present in Chile, their biogeographic affinity with species from the rest of the world has not been studied, with only suggestions of a possible affinity with the Austral/Oceanic entomofauna. On the other hand, the phylogenetic arrangement among the species of the country is not clear, but the history of taxonomic changes suggests a greater kinship between two groups of species according to their similarity in morphological characteristics: 1) C. chilensis, C. ovaticeps and C. spinolae; 2) C. morosus, C. distinguendus and C. hellmichi. To date, no studies have been conducted to directly verify this proposal or to analyze the phylogenetic relationships among Chilean species. In the present study, we evaluated for the first time the biogeographic affinities and phylogenetic relationships of *Camponotus* species present in Chile through the estimation of a global and local phylogeny using molecular data, which were generated for the first time for Chilean species and complemented with sequences obtained from BOLD for species from the rest of the world. Additionally, specimens that could not be identified with the available taxonomic keys and whose morphological characters did not fit the existing species diagnoses were included in the study to evaluate their phylogenetic position and the possibility that they correspond to new species. The results indicate a greater affinity of the Chilean species with the Neotropical and Afrotropical fauna, in disagreement with the hypothesis of a possible relationship with the Austral/Oceanic fauna. Second, the relationships among Chilean species did not follow the expected pattern suggested by the taxonomic history of the species. It seems that certain morphological characteristics evolved independently more than once in different clades of the phylogeny. Finally, profound genetic divergences were found, both among known species and with specimens that could not be identified a priori, which in some cases also extended to lineages, presumably of the same species, suggesting substantial cryptic diversity and the possibility of several undescribed species.

## 2. Introduction

South America has been in contact with different landmasses throughout its history. Approximately 150 million years ago (Ma), it formed part of the supercontinent Gondwana, remaining connected to territories that today correspond to Antarctica, Oceania and Africa (Seton et al., 2012). Around 3 Ma, it came into contact with North America, triggering the biogeographic event known as the “Great American Biotic Interchange” (GABI) (Freitas-Oliveira et al., 2024). As a result, the South American biota exhibits heterogeneous affinities (Segovia & Armesto, 2015; Chávez, 2020), often reflecting relationships with territories to which it is no longer geographically connected.

Within South America, the Chilean territory possesses geographic features that have kept it largely isolated from the rest of the continent for millions of years (e.g., the Andes Mountains, the Pacific Ocean, and the Atacama Desert) (Folguera et al., 2018; Wennrich et al., 2025). The prolonged isolation has resulted in a biota with a high degree of endemism and biogeographic patterns that differ markedly from those of other South American regions (Segovia & Armesto, 2015; Villagrán, 2018). In both flora and fauna, stronger taxonomic affinities with Oceanian components than with Neotropical ones have been reported, reflecting close relationships between the Chilean biota and Gondwana (Boyer & Giribet, 2007; Vera-Escalona et al., 2020; Villagrán & Hinojosa, 2005). In insects, similar patterns have been identified (Arias et al., 2009; Crisci et al., 1991; Palma & Figueroa, 2008), although their specific biogeographic relationships remain poorly known for most taxa. This knowledge gap limits our understanding of how different regions have contributed to the composition of Chilean biodiversity.

Ants represent a well-studied and ecologically important component of terrestrial ecosystems and provide an appropriate model to explore these biogeographic questions. In Chile, the ant fauna comprises 73 described species (Johnson, 2021; Snelling & Hunt, 1975). Research regarding their affinities has been limited or nearly nonexistent, making it difficult to estimate their potential relationships with other regions. This challenge is even greater for species belonging to globally distributed genera, whose affinities may lie across multiple regions. Such is the case of ants in the genus *Camponotus* (Hymenoptera, Formicidae). Camponotus is one of the most diverse ant genera worldwide (>1,000 species) (Bolton, 1995), distributed across the vast majority of the planet’s regions (Fisher & Cover, 2007). Their high species richness, together with pronounced morphological variation associated with worker caste polymorphism, has complicated efforts to infer phylogenetic relationships among species (Mackay & Delsinne, 2009).

For Chilean *Camponotus* species in particular, the literature on potential relationships with the rest of the world is scarce. An early hypothesis was proposed by Kusnezov (1963), who suggested a possible austral affinity between Chilean *Camponotus* species and Oceanian taxa, particularly those from New Zealand. However, this proposal was based largely on the presence of the genus in both regions, which is a weak criterion given its cosmopolitan distribution, and therefore does not exclude affinities with other biogeographic regions where *Camponotus* is also present. As a result of this limited and largely descriptive evidence, the affinities of the Chilean *Camponotus* species with the rest of the world have not yet been formally investigated.

Similarly, the phylogenetic relationships within the Chilean *Camponotus* species have not yet been studied. Within Chile, morphological similarities among certain lineages have historically led to taxonomic confusion, providing a basis to hypothesize potential groups, which are evaluated in this study.

This study addresses two main objectives:

1. To clarify the biogeographic affinities of Chilean *Camponotus* species, particularly testing for a possible austral-Oceanian relationship, and to assess whether their diversity derives from a single evolutionary lineage or from multiple independent origins.
2. To assess the phylogenetic relationships among Chilean species, including hypotheses about potential groups arising from historical taxonomic patterns.

## 3. Materials and Methods

### 3.1. Collection and Storage

Between February and October 2024, *Camponotus* individuals were collected from Arica to Los Lagos, Chile (Table 1). Between 1 and 20 individuals were collected per colony, depending on availability. All individuals from the same colony were pooled in a single container with 95% ethanol and subsequently stored at −20 °C in the Laboratory of Insect Molecular Ecology (LEMIn) at the Institute of Entomology, Universidad Metropolitana de Ciencias de la Educación. Specimens from all known species were included, and when possible, more than one colony per species from geographically distant localities were included to better capture intraspecific variability.

**Table 1.**
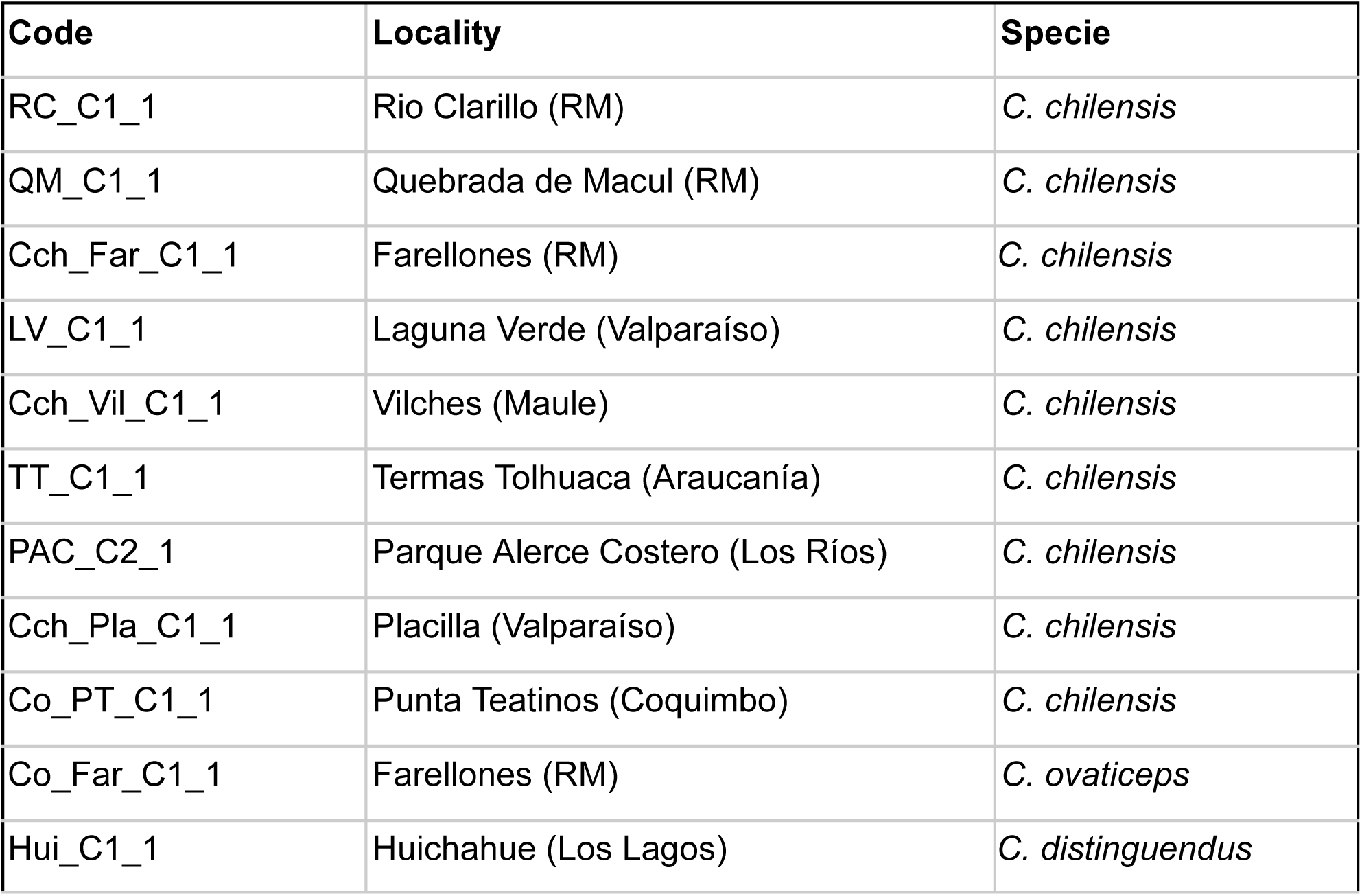
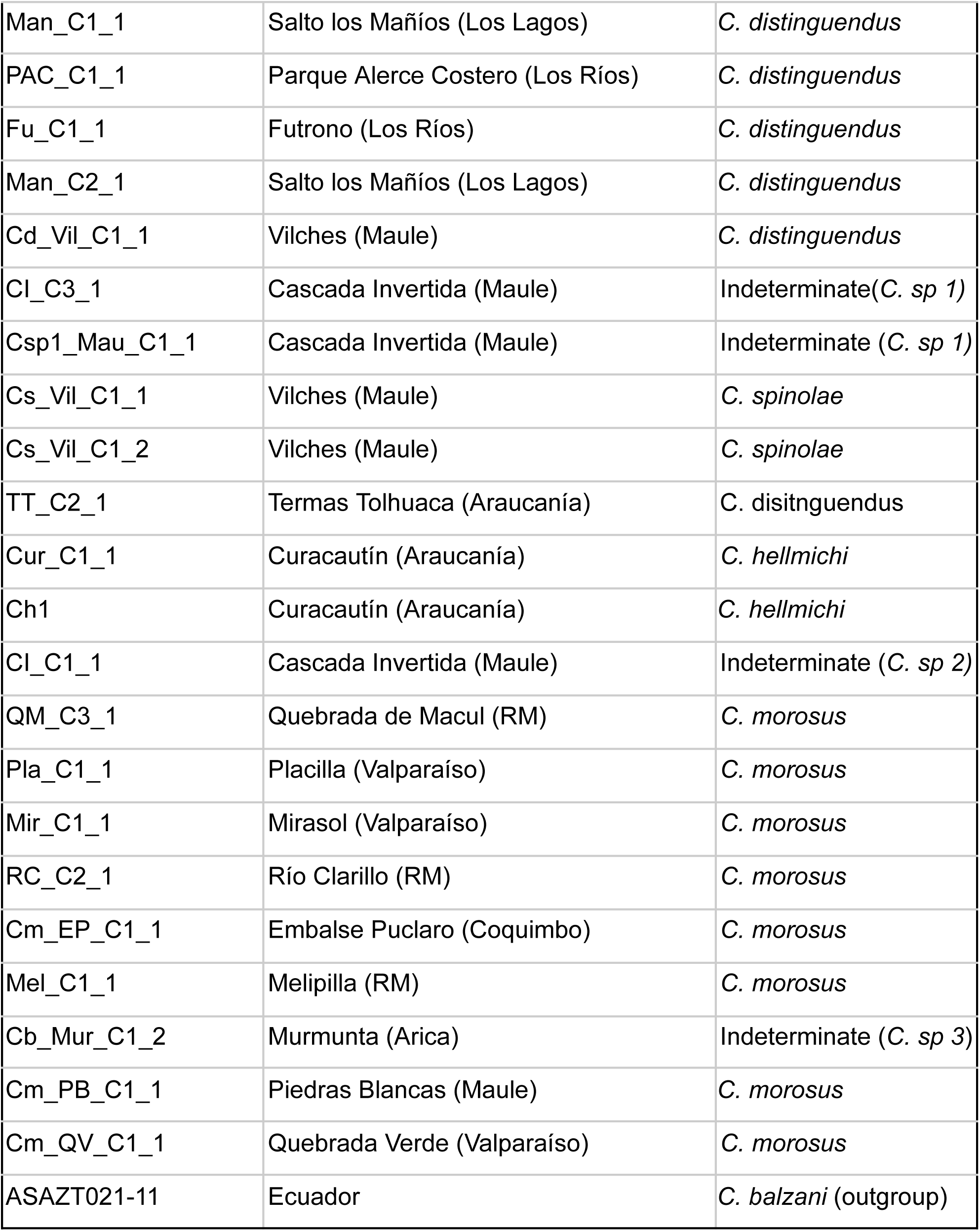
Data associated with Chilean *Camponotus* specimens used in the phylogenetic analyses of this study, including locality, date of collection, and species assignment.

### 3.2. Specimen Identification

Specimens were identified using species descriptions and the dichotomous key provided in Snelling & Hunt (1975). Individuals were assigned to one of the six species currently described for Chile: *Camponotus chilensis* (Spinola, 1851), *Camponotus distinguendus* (Spinola, 1851), *Camponotus ovaticeps* (Spinola, 1851), *Camponotus morosus* (Smith, 1858), *Camponotus hellmichi* Menozzi, 1935, and *Camponotus spinolae* Roger, 1863. In addition, specimens belonging to the genus *Camponotus* but not assignable to any known Chilean species were included in order to evaluate whether these individuals correspond to phenotypic variants of known species or to potentially undescribed taxa.

#### 3.2.1. Taxonomic Background

Due to morphological similarity, several species have historically been treated as varieties of others:

On the one hand, *C. ovaticeps* and *C. spinolae* were both historically considered varieties/subspecies of *C. chilensis*: *C. ovaticeps* was treated as *C. chilensis var. ovaticeps* (Emery, 1894; Roger, 1863), and *C. spinolae* was treated as *C. chilensis var. rufficornis* by Emery (1894) and Forel (1907). These three species share the presence of dense golden pilosity on the dorsal portion of the gaster, which obscures visibility of the underlying integument.

On the other hand, *C. morosus* was treated as *C. distinguendus morosus* by Emery (1894), while *C. hellmichi* was originally described as *C. morosus var. hellmichi* by Menozzi (1935). These three species share whitish and sparse pilosity on the dorsal portion of the gaster, allowing the integument to remain visible.

Based on this, two groups of closely related species can be hypothesized:

1. a group composed of *C. chilensis*, *C. ovaticeps*, and *C. spinolae* (hereafter referred to as the “chilensis group”), and
2. a group composed of *C. distinguendus*, *C. morosus*, and *C. hellmichi* (hereafter referred to as the “distinguendus group”).

### 3.3. DNA Extraction and Amplification

DNA extraction was performed using the DNeasy Blood and Tissue Kit (Qiagen, Germany), following the manufacturer’s protocol. For each colony, the largest and best-preserved individual was selected, from which the three legs on the right side were removed. To enhance extraction efficiency, legs were cut into small pieces to maximize exposure of muscle tissue and improve lysis.

Once extracted, DNA was used to amplify the mitochondrial gene Cytochrome Oxidase I (subunit 1) (COI) by PCR, using LCO and HCO primers from Folmer (1994). The reaction mixture (total volume 30 µL) included: Go-Taq (0.4 µL), buffer with loading dye for electrophoresis (6 µL), BSA (0.3 µL), dNTPs (0.6 µL), MgCl₂ (2.4 µL), HCO and LCO primers (1.5 µL each), ultrapure water (16.3 µL), and DNA template (1 µL). The thermal cycling protocol consisted of: 95 °C for 1 min (“hot-start”), followed by 5 cycles of denaturation (94 °C, 30 s), annealing (48 °C, 40 s), and extension (72 °C, 1 min). This was followed by 30 cycles of denaturation (94 °C, 30 s), annealing (52 °C, 40 s), and extension (72 °C, 1 min), with a final hold at 4 °C. PCR success was verified by electrophoresis on 1% agarose gel stained with SYBR Safe (Thermo Fisher Scientific Inc., USA). Samples that amplified successfully were stored at −20 °C until sequencing at MACROGEN (Santiago, Chile).

### 3.4. Sequence Editing and Phylogenetic Analyses

Sequences were visualized and edited using BioEdit v7.7.1 (Hall, 1999) and CodonCode Aligner (v11.0.2, CodonCode Corporation, Tokyo, Japan). Phylogenetic reconstructions were carried out at two spatial scales:

1. a global *Camponotus* phylogeny to evaluate the monophyly of Chilean species and their biogeographic affinities, and
2. a local phylogeny to assess relationships among Chilean species.

For the global phylogeny, in addition to Chilean sequences generated in this study, all *Camponotus* COI sequences available in the Barcode of Life Data System (BOLD) (Ratnasingham & Hebert, 2007) were downloaded in .TSV format. Sequences were filtered to retain only those identified at species level, and one high-quality sequence per species was selected. This reduced the dataset from 23,504 sequences to 322. The phylogeny was inferred using Maximum Likelihood in RAxMLGUI v2.0.10 (Edler et al., 2021), first determining the best-fit evolutionary model (GTRGAMMA) and then conducting a search for the best tree with 1,000 rapid bootstrap replicates. The resulting tree was visualized and rooted in FigTree v1.4.4 (Rambaut, 2018), using as outgroup species currently placed in the genera *Colobopsis* and *Dinomyrmex*(Ward et al., 2016), previously considered part of *Camponotus*. To represent species’ biogeographic origins, each was classified according to the regions proposed by Morrone (2015b), and visualization was performed with Iroki Phylogenetic Tree Viewer (Moore et al., 2020).

The local phylogeny (Chilean species only) was also inferred using Maximum Likelihood in RAxMLGUI v2.0.10. The best-fit evolutionary model was GTRGAMMAX. The final analysis included 1,000 rapid bootstrap replicates prior to the search for the best tree. The tree was visualized and rooted in FigTree v1.4.4 using *Camponotus balzani* as outgroup, based on results from the global phylogeny.

To evaluate genetic divergence, pairwise genetic distances were calculated using the Kimura 2-Parameter (K2P) model (Kimura, 1980) in MEGA v11.0.13 (Tamura et al., 2021). For this analysis, up to two specimens per subclade and/or species were included whenever possible.

### 3.5. Species Distribution Mapping

Distribution maps were generated using occurrence data from field collections, georeferenced records obtained from the open-source biodiversity platform GBIF, locality records reported in taxonomic catalogs (Snelling & Hunt, 1975), and specimens from the collection of the Institute of Entomology at the Universidad Metropolitana de Ciencias de la Educación (IE-UMCE). In the case of GBIF, records with clearly out-of-range distributions were reviewed: photographic vouchers were inspected to verify correct species identification, and records lacking photographs were excluded.

Maps were produced in QGIS Long Term Release version 3.40.11 (QGIS Development Team, 2024). Geographic layers “countries,” “oceans,” and “lakes” were downloaded from Natural Earth (Kelso & Patterson, 2010), and the “Regions of Chile” layer was obtained from the map repository of the Chilean National Congress Library (BCN).

## 4. Results

### 4.1. Global Phylogeny

The global phylogeny of the genus *Camponotus* (Figure 1) shows that the species present in Chile form a monophyletic group, with moderate bootstrap support (50 to 75). This clade has *Camponotus balzani*, a Neotropical species, as its sister species. Within the clade it is possible to observe the existence of two subclades, one with high support (75 to 100) that includes the great majority of species, and one with low support that includes specimens identified as C. morosus and C. sp3.

**Figure 1:**
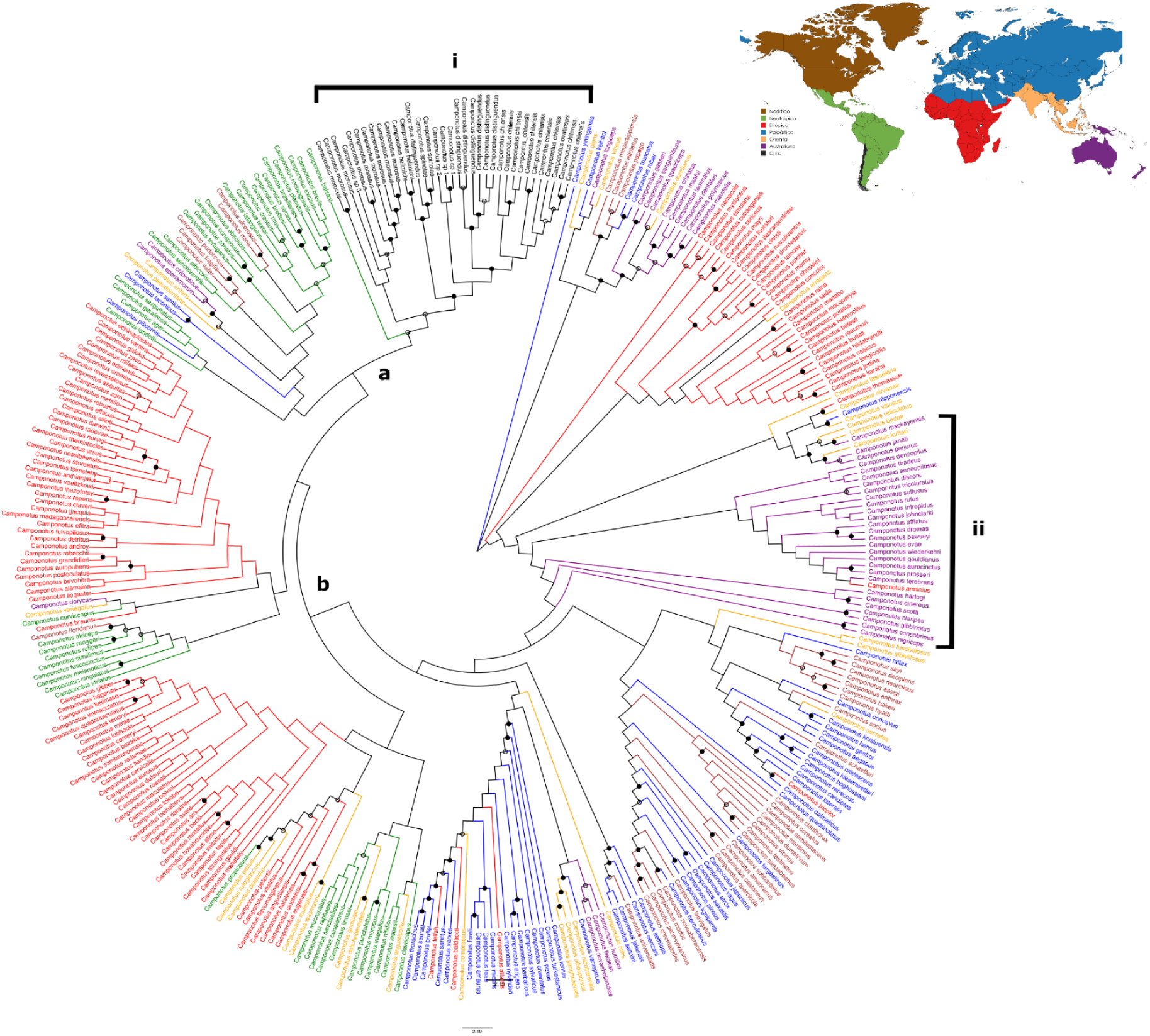
Global phylogeny of *Camponotus* species inferred from COI sequences representing 328 species. Each species and branch is colored according to its biogeographic origin: Neotropical (green), Nearctic (brown), Palearctic (blue), Ethiopian (red), Oriental (orange), Oceanian (purple), and Chile (black). The Chilean clade is marked as (i). Clades (a) and (b), and sector (ii) are referenced in the Results and Discussion. The phylogeny is rooted using species currently placed in the sister genera *Colobopsis* and *Dinomyrmex* (Ward et al., 2016). Bootstrap values are shown as circles at each node: filled circle = 75–100; empty circle = 50–75; no circle = 0–50. For detailed bootstrap values, see Figure S1 (Appendix).

Clade (a), which contains 40 species (including the Chilean species already described), is mostly composed of Neotropical species (70%) and, to a lesser extent, Nearctic species (12.5%), Palearctic (7.5%), Oriental (5%), and Oceanian species (5%). In turn, this clade forms part of a larger clade (b), dominated mostly by Afrotropical species.

Most species with Oceanian distributions are found in early diverging clades (ii), and the rest are dispersed throughout the phylogeny. In general, the deeper clades show low bootstrap support, whereas the more recent clades present high support values, indicating greater phylogenetic resolution in the terminal relationships.

### 4.2. Local Phylogeny (Chile)

Regarding the specimens of already described species: In the phylogeny of the Chilean species of *Camponotus* (Figure 2), several clades can be distinguished. Clade “a” groups the species *Camponotus chilensis*, *Camponotus ovaticeps* and *Camponotus distinguendus*, with genetic distances that range from 0 to 5.7% among their populations (Table S1) and with high bootstrap support (50–75). Clade “b” groups the species C. spinolae, C. hellmichi and *Camponotus* sp4, with genetic distances that range between 0.1 and 5.1% among their populations (Table S1) and with high bootstrap support (75–100). Finally, clade (c) includes C. morosus, with genetic distances that vary between 0.3% and 9.3% among their populations, and with intermediate bootstrap support (50–75). The genetic distances between clades are considerably greater: between clades “a” and “b” they range between 8.4% and 11.1%; between clades “a” and “c”, between 13.6% and 18.8%; and between clades “b” and “c”, between 11.4% and 15.5%.

**Figure 2:**
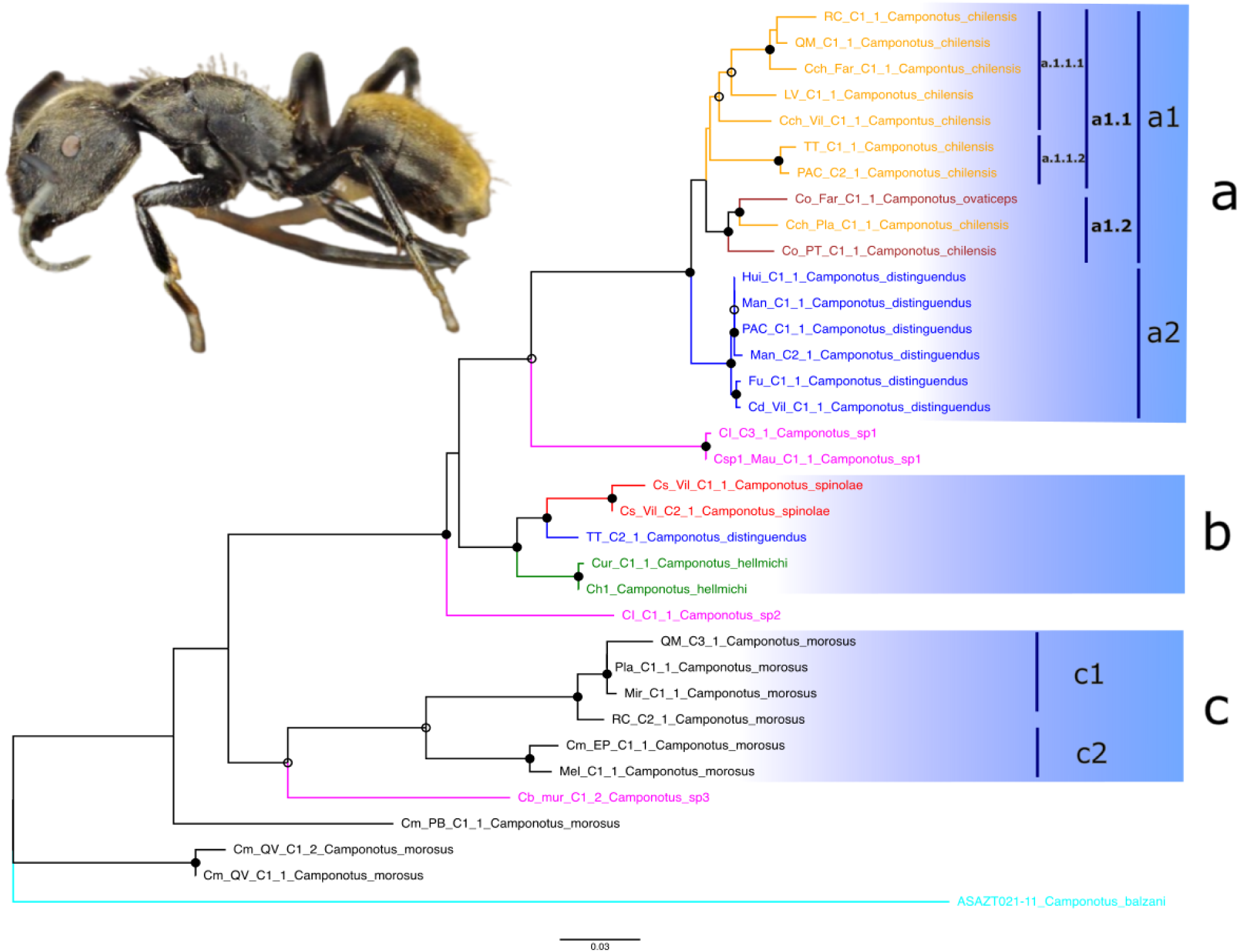
Phylogeny of Chilean *Camponotus* based on COI (subunit 1). Thirty-five sequences were used, including *C. balzani*(cyan) as outgroup. Species colors: *C. chilensis* (orange), *C. ovaticeps* (brown), *C. spinolae* (red), *C. distinguendus* (blue), *C. morosus* (black), *C. hellmichi* (green), *C. sp.* (unidentified lineages; fuchsia). Clade a includes *C. chilensis*, *C. ovaticeps*, *C. distinguendus*; clade b includes *C. hellmichi*, *C. spinolae*, *C. sp4*; clade c includes *C. morosus*. Bootstrap values: filled circle = 75–100; empty = 50–75; absent = 0–50. For detailed values, see Figure S2 (Appendix).

Clade “a” subdivides into 2 subclades, “a1” and “a2”, with high bootstrap support. Subclade “a2” groups the great majority of populations of *Camponotus distinguendus* with internal genetic distances between 0 and 0.7%. The other subclade “a1” groups the populations of C. chilensis and C. ovaticeps, with genetic distances between 0.8 and 5.7%. This subclade in turn subdivides into 2 subclades, “a1.1” and “a1.2”, with low bootstrap support (0 to 50). “a1.1” groups the great majority of C. chilensis populations with internal distances between 0.8 and 5.1%, whereas “a1.2” groups the two populations of C. ovaticeps and the population Cch_Pla_C1_1 of C. chilensis, with internal distances between 2.7 and 3.4%. Finally, clade “a1.1” subdivides into 2 subclades, “a1.1.1” and “a1.1.2”, both with low bootstrap support. “a1.1.1” groups the C. chilensis populations of the south–central part of the country, with internal genetic distances between 0.9 and 3.6%, whereas “a1.1.2” groups the two southernmost C. chilensis populations with a genetic distance of 0.8%.

It is also important to mention that the maximum genetic difference among populations identified as C. chilensis is 5.7%, which is higher than that found among some populations of C. distinguendus. In the case of the specimen identified as C. ovaticeps, it shows a difference of 2.7% compared to the C. chilensis population Cch_Pla_C1_1, a lower percentage difference than that between other C. chilensis populations.

Clade “b” is composed of the specimens of the species C. spinolae, C. hellmichi and the specimen TT_C2_1 categorized as C. distinguendus. The internal genetic distances of the clade range between 0.1 and 5.1%. Within the clade, the specimens of the species C. spinolae and C. hellmichi occur in separate subclades. In the case of specimen TT_C2_1, it shows a minimum genetic difference of 3.1% from the rest of the clade.

Clade “c” groups most individuals assigned to C. morosus, with internal genetic distances between 0.3 and 9.3%. This clade subdivides into 2 subclades: “c1” and “c2”. These show a minimum genetic distance of 7.4% between them, and internal distances of 0.3 to 3.2% for “c1”, and 1.8% between the two specimens of “c2”.

Finally, outside the clade containing all the other sequences is Cm_PB_C1_1, and outside this, the specimens Cm_QV_C1_1 and Cm_QV_C1_2, all of them preliminarily assigned to C. morosus. Cm_PB_C1_1 showed a minimum genetic difference of 11.8% relative to the other samples, and the samples from the Cm_QV population showed a minimum difference of 11.4%. These populations did not show relevant morphological differences (Figure S4, Appendix) that would justify their classification as a different species according to the taxonomic key used (Snelling & Hunt, 1975).

Regarding minimum genetic distances between species (Table 2), all were greater than 3 percent except between C. chilensis and C. ovaticeps, with a value of 2.7%. Regarding internal genetic distances per species (Table 2), and considering the species identity threshold for ants of 2–3% (Smith et al., 2005): in terms of minimum distance, only C. ovaticeps showed a value greater than 2%. In terms of maximum distance, C. distinguendus, C. chilensis, C. ovaticeps and C. morosus all showed distances greater than 3%.

**Table 2:** Minimum genetic distances between species and internal genetic distances per species. Nomenclature: Cd: C. distinguandus; Cch: C. chilensis; Co: C. ovaticeps; Ch: C. hellmichi; Cs: C. spinolae; Cm: C. morosus. Percentage expressed as a proportional value (i.e., 1=100%). For more details on genetic distances between samples, see Table S1 (Appendix).

| Minimum genetic distance between species |  |  |  |  |  |  |
| --- | --- | --- | --- | --- | --- | --- |
| <u>Species</u> | <u>C.d</u> | C.ch | <u>C.o.</u> | <u>C.h</u> | <u>C.s</u> | <u>C.m</u> |
| **C.d** | - | - | - | - | - | - |
| **C.ch** | 0.034 | - | - | - | - | - |
| **C.o.** | 0.034 | 0.027 | - | - | - | - |
| **C.h** | 0.039 | 0.091 | 0.084 | - | - | - |
| **C.s** | 0.031 | 0.095 | 0.091 | 0.043 | - | - |
| **C.m** | 0.124 | 0.128 | 0.12 | 0.114 | 0.117 | - |
| Internal genetic distance by species |  |  |  |  |  |  |
| <u>Species</u> | <u>Minimum</u> |  | <u>Maximum</u> |  |  |  |
| **C.d** | 0 |  | 0.1 |  |  |  |
| **C.ch** | 0.008 |  | 0.057 |  |  |  |
| **C.o.** | 0.034 |  | 0.034 |  |  |  |
| **C.h** | 0.001 |  | 0.001 |  |  |  |
| **C.s** | 0.011 |  | 0.011 |  |  |  |
| **C.m** | 0.003 |  | 0.151 |  |  |  |

Regarding the specimens classified as not belonging to any described species: None of the samples nested within any known species, and all showed high genetic differentiation relative to their closest phylogenetic relatives. Minimum genetic distances were: *C. sp. 1*: 8.1%; *C. sp. 2*: 7.5%; and *C. sp. 3*: 10.9%.

## 5. Discussion

### 5.1. Biogeographic affinity of Chilean species

When observing the global phylogeny, it is possible to see that the clade grouping the Chilean species (i) is distant from the clade that contains most Australian species (ii), suggesting a distant relationship between these two groups. This information goes against the hypothesis proposed by Kusnezov (1963) of a biogeographic affinity between the Chilean entomological fauna and the Austral–Oceanian fauna, and instead suggests the existence of a stronger affinity with other biogeographic areas.

Considering South America’s recent biogeographic history, another possible biogeographic affinity of the Chilean clade could be with the Nearctic group. In this case, the ancestor of the Chilean group and its sister species *C. balzani* could have crossed (from or toward North America) through the Isthmus of Panama during the Great American Biotic Interchange about 3 million years ago (Chávez, 2020). However, this proposal faces several difficulties. First, the geographic barriers that currently prevent faunal exchange between Chile and the rest of South America were already fully present and established by that time (Boschman, 2021; Hartley & Chong, 2002), meaning that migration would also have been strongly restricted then, especially in the central and northern parts of the country, as remains the case for many species today (Kellner et al., 2013; Nunes, 2017). Second, there is no substantial presence of Nearctic species closely related to the Chilean clade, as would be expected under this scenario. Instead, a greater number of Afrotropical species are present. Thus, the possibility of a Nearctic affinity and potential Nearctic origin for the Chilean species is weakly supported.

The presence of Afrotropical species closely related to the Chilean clade suggests that the *Camponotus* species present in Chile and much of the Neotropics may have an ancient origin dating back to the time of Gondwana. This biogeographic affinity with African elements has been reported in other cases, including both fauna and flora (Bond et al., 2015; Cidade et al., 2019; Moreira-Muñoz, 2011). In Chile, examples reporting this relationship are more limited and particularly scarce for entomofauna (Martins-Neto et al., 2003; Ragionieri et al., 2023), where relationships more commonly reflect Gondwanan/Austral affinities among different insect clades (Arias et al., 2009; Crisci et al., 1991; Hodgson & Miller, 2002; Morrone, 2006; Palma & Figueroa, 2008; Szwedo, 2004).

South America shows the lowest proportion of biogeographic studies among inhabited continents (Beheregaray, 2008), despite paradoxically being considered the territory with the highest biodiversity worldwide. This highlights the need for further research to better understand the biogeographic relationships of the continent’s biota. The results of this study contribute to improving understanding of the composition of South American entomofauna, particularly the myrmecological fauna of Chile and the Andean region, suggesting that Chilean *Camponotus* species have a closer relationship with Afrotropical and Oriental fauna, in contrast to other insects whose affinities appear to be more closely related to the Oceanian fauna.

### 5.2. Relationships among Chilean species and phylogenetic value of the main diagnostic morphological characters

Considering the history of taxonomic changes within the genus *Camponotus* in Chile and the morphological similarity regarding the absence/presence of pilosity on the gastral tergite, this study hypothesized the existence of two groups with greater phylogenetic proximity (“chilensis group” and “distinguendus group”). When observing the results of the local phylogeny (Figure 2), it becomes evident that the phylogenetic proximity suggested by taxonomic history is not consistent with the ancestor–descendant pattern observed in the tree. On the contrary, this phylogeny reveals relationships that differ greatly from the two hypothesized groups. In addition to gastral pilosity, among the most commonly used characters in species descriptions and dichotomous keys for Chilean *Camponotus* are the presence/absence of lateral cephalic pilosity and variation in antennal coloration. Although these characters remain taxonomically useful, the results of this study also suggest a limited phylogenetic value for these traits among Chilean species (Figure 6).

### 5.3. Molecular support for the described species

Of the six species described to date, all exhibit genetic distances greater than 2–3% from one another, exceeding the species identity threshold proposed for ants (Smith et al., 2005). The only case in which this distance falls between 2–3% is between the specimen Co_Far_C1_1, identified as *C. ovaticeps*, and the population Cch_Pla_C1_1 of *C. chilensis*, which show a difference of 2.7%. Despite this, their marked morphological differences regarding cephalic and thoracic pilosity, along with the proximity of their collection sites (∼110 km), support their validity as distinct taxa. Additionally, the entire distribution of *C. ovaticeps* completely overlaps with that of *C. chilensis* (Figures 3 and 5), with both species occurring in sympatry, which further supports their status as distinct species.

**Figure 3:**
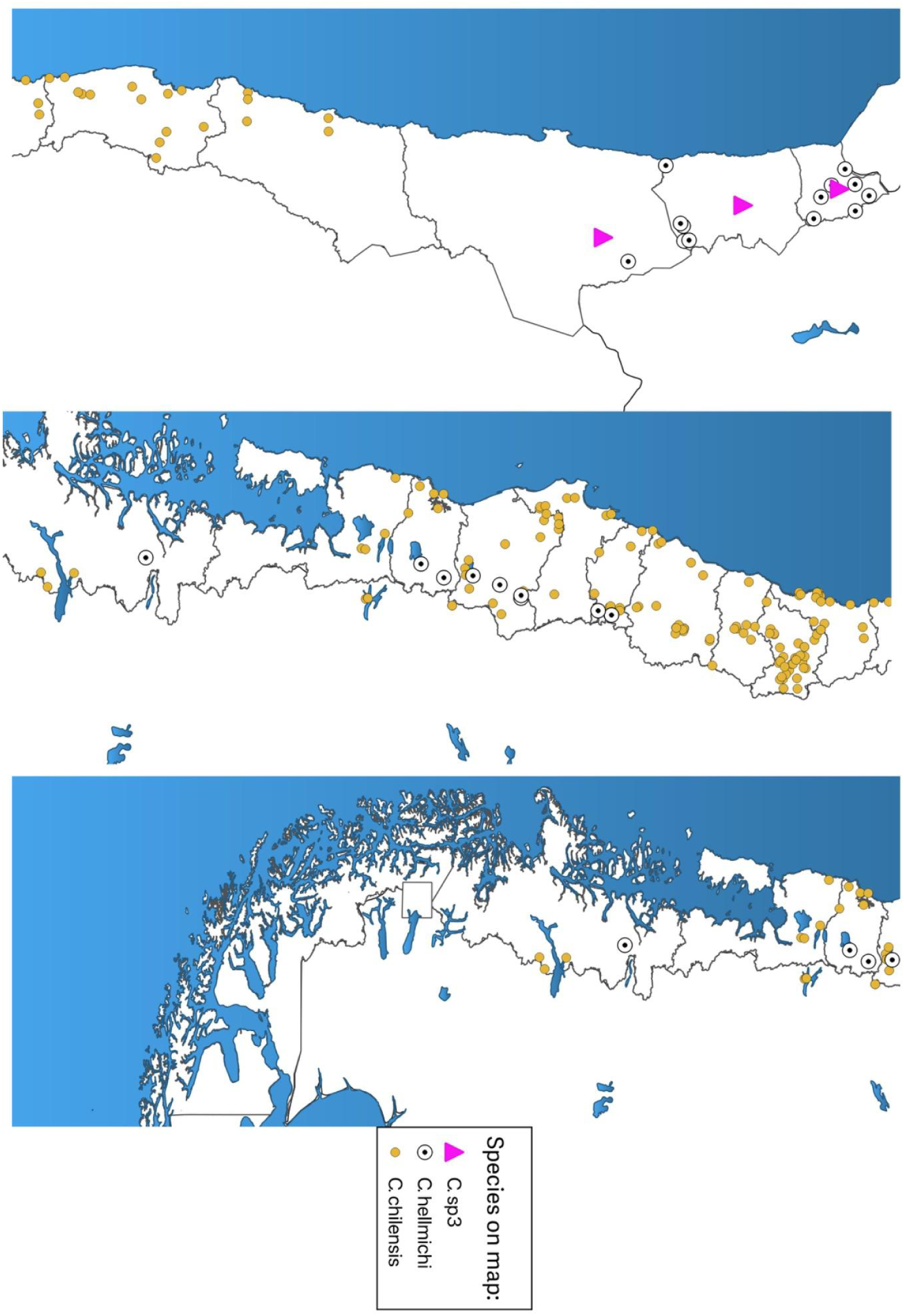
Map showing distribution points of the species *Camponotus chilensis, Camponotus hellmichi,* and *Camponotus sp3.* Maps generated using the QGIS program (2024).

**Figure 4:**
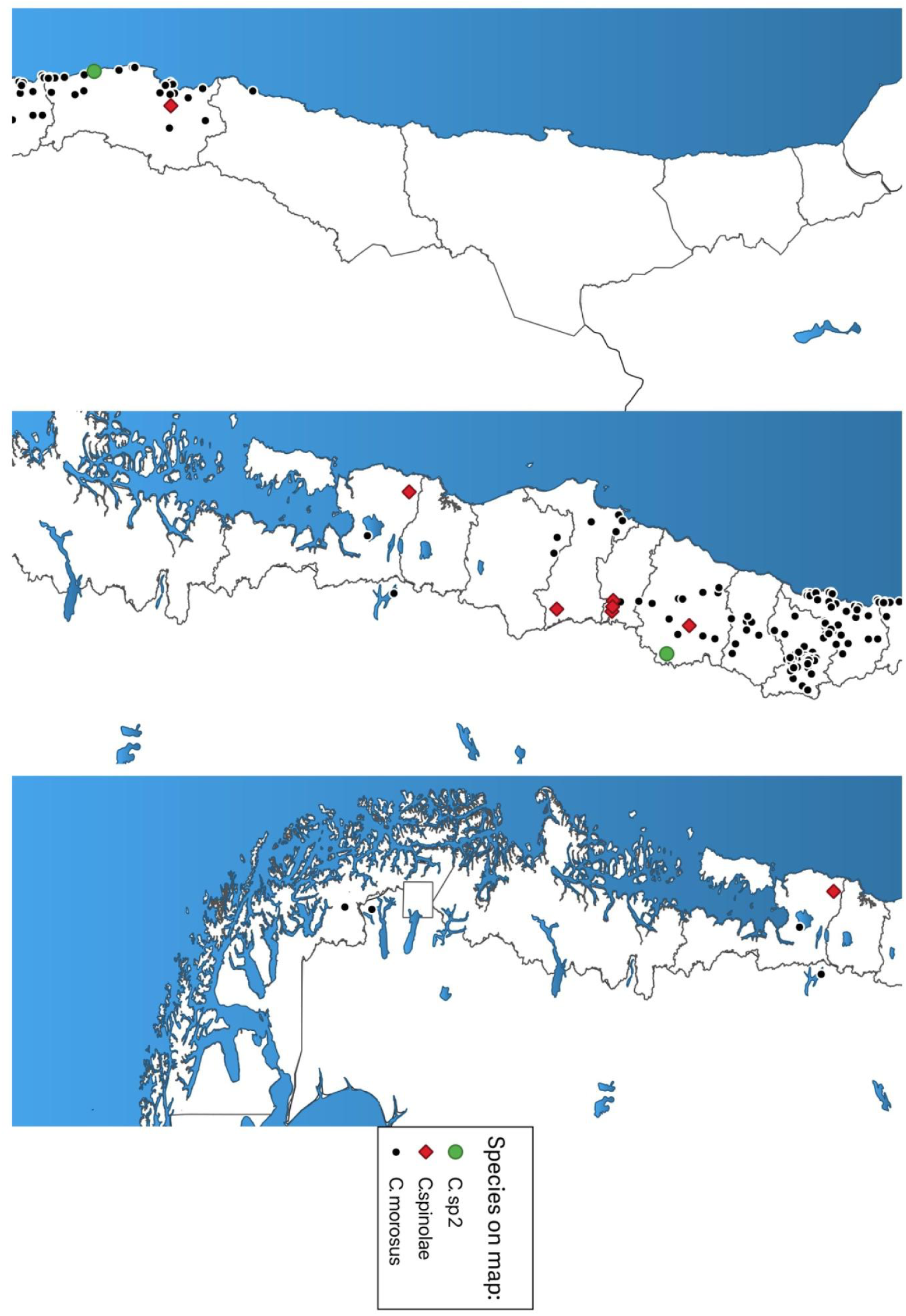
Map showing distribution points of the species *Camponotus spinolae*, *Camponotus morosus*, and *Camponotus sp2*. Maps generated using the QGIS program (2024).

**Figure 5:**
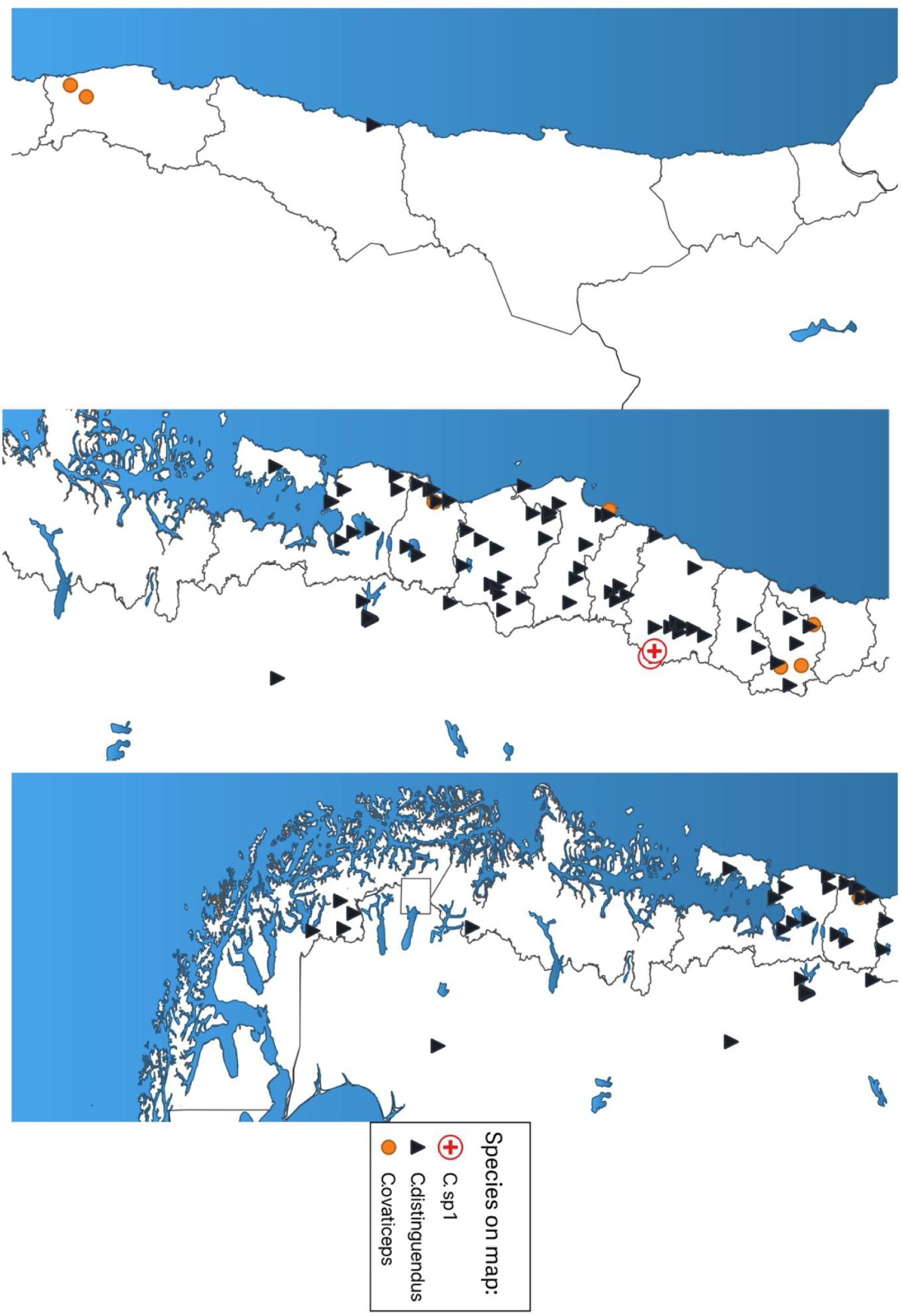
Map showing distribution points of the species *Camponotus distinguendus*, *Camponotus ovaticeps*, and *Camponotus sp1*. Maps generated using the QGIS program (2024).

**Figure 6:**
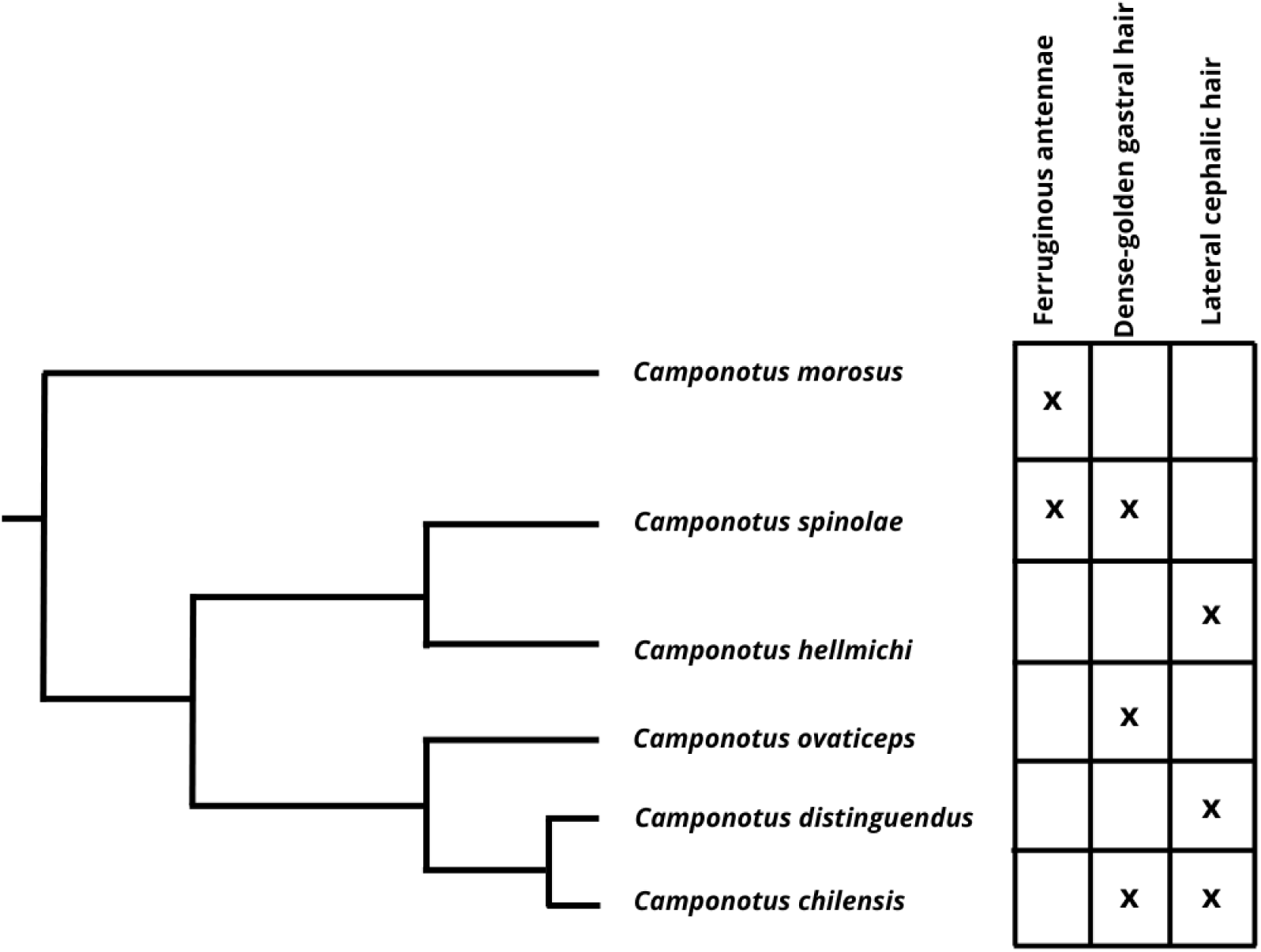
Phylogenetic distribution of some of the main morphological characters of the *Camponotus* species present in Chile. The three characters with the greatest relevance for distinguishing species according to the key of Snelling and Hunt (1973) were considered: red antennae, dense/golden gastral pilosity, and lateral cephalic pilosity. The specimen *Camponotus* sp. 4 is included to indicate that the character “lateral cephalic pilosity” is not exclusive to the *chilensis–distinguendus* clade.

Despite the points mentioned above, it is important to highlight the morphological similarity between *C. ovaticeps* and *C. chilensis*, whose main distinguishing traits are the absence/low abundance of lateral cephalic pilosity and a relatively lower abundance of golden gastral pilosity in *C. ovaticeps* (Spinola, 1851). During field sampling across the country, a gradient was found in the abundance of lateral pilosity and dorsal gastral pilosity, suggesting that this character may not be ideal for distinguishing between the two species.

### 5.4. Intraspecific variability

Regarding individuals identified as *C. chilensis* using the key of Snelling & Hunt (1975): in many cases, high genetic distances were found between individuals or groups of individuals, reaching values greater than 2% or even 5%. Such differences would be expected among populations separated by large geographic distances, which could lead to isolation and divergence over thousands of years (Relethford, 2004). However, in this study, many of the highly divergent populations are separated by only tens of kilometers. For example, RC_C1_1 and Cch_Far_C1_1 show a genetic distance of 2.4% and are separated by roughly 50 km. A more extreme case is that of Cch_Pla_C1_1 and LV_C1_1, with a genetic distance of 4.3% and collection sites less than 15 km apart. As mentioned earlier, this genetic variability is accompanied by morphological variability in the abundance of lateral cephalic pilosity and gastral pilosity. This combination of high genetic divergence and morphological variability at short geographic distances suggests that specimens currently classified as *Camponotus chilensis* may not correspond to a single species, but rather to a species complex with high morphological similarity, complicating differentiation.

In contrast, most individuals identified as *C. distinguendus* show low genetic distances (<0.7%), even between populations more than 500 km apart (e.g., Cd_Vil_C1_1 and Fu_C1_1). The only exceptional case within this species is TT_C2_1, present in the phylogeny in clade (b), which, despite morphological similarity, has a surprising distance of 10% from the rest of the *C. distinguendus* populations. Additionally, it belongs to a completely different clade with high support (clade “b”, Figure 2), showing greater phylogenetic affinity with *C. spinolae* and *C. hellmichi*.

In the case of specimens identified as *C. ovaticeps*, it is relevant to mention that Co_Far_C1_1 was classified as this species following the available key (Snelling & Hunt, 1975), due to the presence of golden gastral pilosity, black antennae, and absence of lateral cephalic pilosity. However, the original description (Spinola, 1851) does not mention several morphological traits characteristic of this taxon, such as the ferruginous coloration of the legs present in all female castes. This raises doubts about whether this taxon truly belongs to this species, casting uncertainty on the usefulness of the available key for effectively distinguishing among the lineages present in the country. Thus, it is possible that this specimen represents a novel taxon within clade “a” rather than belonging to *C. ovaticeps*.

Regarding individuals identified as *C. morosus* using the key of Snelling & Hunt (1975): as with *C. chilensis*, in many cases there were high genetic distances among individuals or groups of individuals, but in *C. morosus* these differences were even larger, reaching 7% between subclades “c1” and “c2” within clade “c” (Figure 2), and more than 11% with individuals Cm_PB_C1_1, Cm_QV_C1_1 and Cm_QV_C1_2. Unlike *C. chilensis*, *C. morosus* specimens displayed very low or no morphological differences (Figure S4, Appendix), suggesting the existence of cryptic species within this group of specimens.

The appearance of similar morphologies in different branches of the phylogeny, both in *C. chilensis*, *C. distinguendus*, and *C. morosus*, is not unusual in the study of the genus *Camponotus*. Several cases exist where species possess similar traits despite being distantly related phylogenetically, or even appearing as the most genetically distant species (Ki-Gyong & Byung-Jin, 2006; McArthur & Leys, 2006). Such morphological similarity extends beyond species within the genus, occurring also within and outside the tribe Camponotini. For example, in the sister genus *Colobopsis*, morphological similarity is so pronounced among species that relationships among some species groups have only been resolved using molecular tools (Ward et al., 2016).

### 5.4. Evidence of new taxa and their implications for the diversity of the genus *Camponotus* in Chile

Three specimens that could not be identified as any known species due to their morphology (*C. sp1*, *C. sp2* and *C. sp3*) were included in the phylogenetic analysis. All showed differences greater than 3% and did not nest within any already described species. This supports the hypothesis that these individuals represent novel taxa within the genus in Chile.

Regarding the distribution of *C. sp3*, the IE-UMCE collection contains specimens with similar characteristics that were not identified, and some of them had been catalogued as *C. hellmichi*. We believe this erroneous classification is due to similarity in some morphological traits used in the identification keys (absence of lateral cephalic pilosity, black antennae, and absence of golden gastral pilosity), but other traits—such as the smooth, shiny integument in *C. sp3*versus the punctate, opaque integument in *C. hellmichi*, allow easy differentiation (Figure S5, Appendix). Furthermore, the distribution of *C. sp3* matches the northern distribution points of *C. hellmichi* noted in Snelling & Hunt (1975) (Figure 3), suggesting that this confusion may have occurred in that study as well. This implies that the actual distribution of *C. hellmichi* may be much more restricted, closer to its type locality (Villarica Volcano) in southern Chile (Menozzi, 1963).

The three unidentified specimens are found entirely or partly in the Andean region of northern and south-central Chile (Figures 3, 4 and 5), areas adjacent to the borders with Bolivia and Argentina. Therefore, it is important to consider and investigate whether these specimens may correspond to taxa already described for those neighboring countries but not yet recognized in Chile.

Based on what has been discussed, it is reasonable to propose that the material of *Camponotus* from Chile analyzed in this study includes species that require description and species not yet recognized for the country. Some of these may be cryptic species currently considered part of known taxa due to morphological similarity, while others show clear morphological and genetic differences. Overall, the diversity of the genus *Camponotus* in Chile appears to be considerably underestimated, with a potential number of species much greater than currently acknowledged.

### 5.5. Limitations of the research and future challenges

The use of the “COI subunit 1” sequence to estimate phylogenetic relationships is a widely applied methodology in the study of evolutionary relationships across diverse organisms (Andrade-Souza et al., 2017; Trontelj et al., 2005; Vandewoestijne et al., 2004). To estimate the biogeographic affinities of *Camponotus* species in Chile, this study benefited greatly from the large number of COI sequences available in public databases, which allowed a first approximation of phylogenetic relationships among hundreds of species representing all world regions. However, some parts of the global phylogeny show low support, particularly deeper clades or older nodes. Therefore, this estimation should be interpreted as a preliminary hypothesis regarding relationships among *Camponotus* species worldwide and not as a definitive conclusion. Future research would benefit from including a greater number of genetic regions and species to generate a more robust tree that helps clarify relationships within this highly diverse genus. This represents a major challenge requiring collaboration among researchers worldwide, including as much diversity within the genus as possible.

On the other hand, the number of available Neotropical sequences (49) is very low relative to the large number of described species in the region (AntWeb, 2024), which may strongly affect the evaluation of Chilean monophyly. Including more species in future analyses—particularly from countries adjacent to Chile (Argentina, Bolivia, and Peru), would be crucial for rigorously confirming or rejecting monophyly of the Chilean clade.

Regarding the *Camponotus* species present in Chile: first, the low correlation between morphological similarity and phylogenetic proximity presents a challenge in understanding the processes underlying these patterns. Studies are needed to investigate the evolutionary and ecological mechanisms that have shaped the phenotypes observed today, which vary greatly even among closely related lineages, and conversely appear similar among lineages with deep divergences. Second, it is necessary to formally describe the new taxa that show high genetic differentiation (>2–3%), with or without accompanying morphological differences, in order to resolve the true species diversity present in the country. Alongside this, new taxonomic keys should be developed to effectively distinguish among lineages based on their morphological differences.

Finally, to increase the robustness and representativeness of each species, future studies should incorporate more samples from taxa with low representation, such as *C. ovaticeps*, *C. hellmichi*, *C. spinolae*, *C. sp1*, *C. sp2*, *C. sp3*, and certain individuals of *C. chilensis* and *C. morosus* that appeared genetically distant.

## Acknowledgements

The authors thank Mario Elgueta and Maribel Beltrán for their kind assistance, and access to the collections of the Museo Nacional de Historia Natural (MNHN) and the Insect Collection of the Instituto de Entomología, Universidad Metropolitana de Ciencias de la Educación (UMCE-IE), respectively. This work was partially supported by projects DIUMCE-08−2025−ID and FONDECYT (11220706). BA thanks the support from a Beca de Magister ANID.

## Declarations

Conflict of interest. All the authors declare no conflict of interest.

## Appendix

**Table S1:**
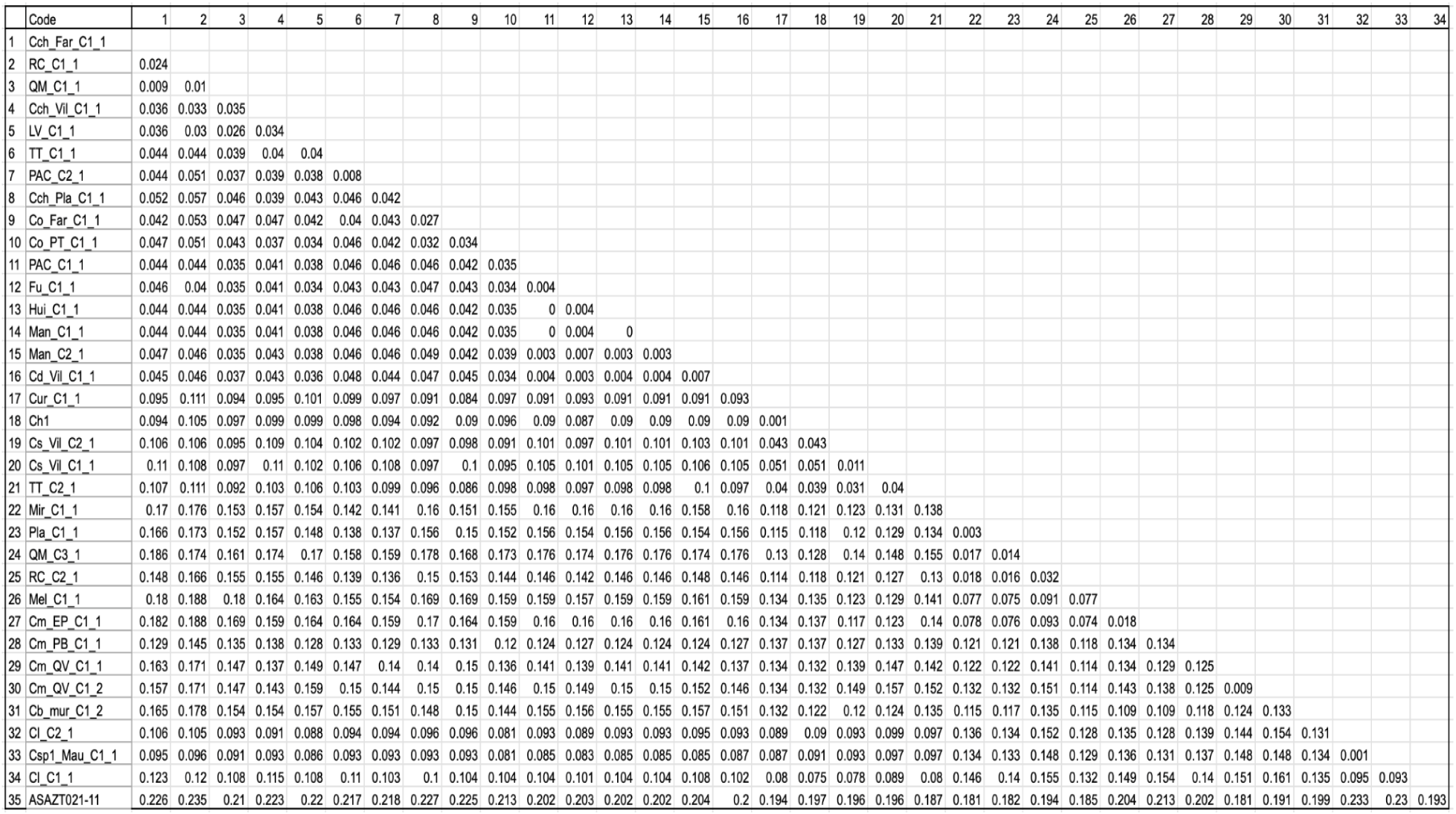
Table of genetic distances (K2P; Kimura, 1980) among *Camponotus* lineages present in Chile. For this calculation, the COI gene (subunit 1) was used, and distances were calculated in the MEGA 11 program (Tamura et al., 2021). Distances were truncated to the third decimal place.

**Figure S1:**
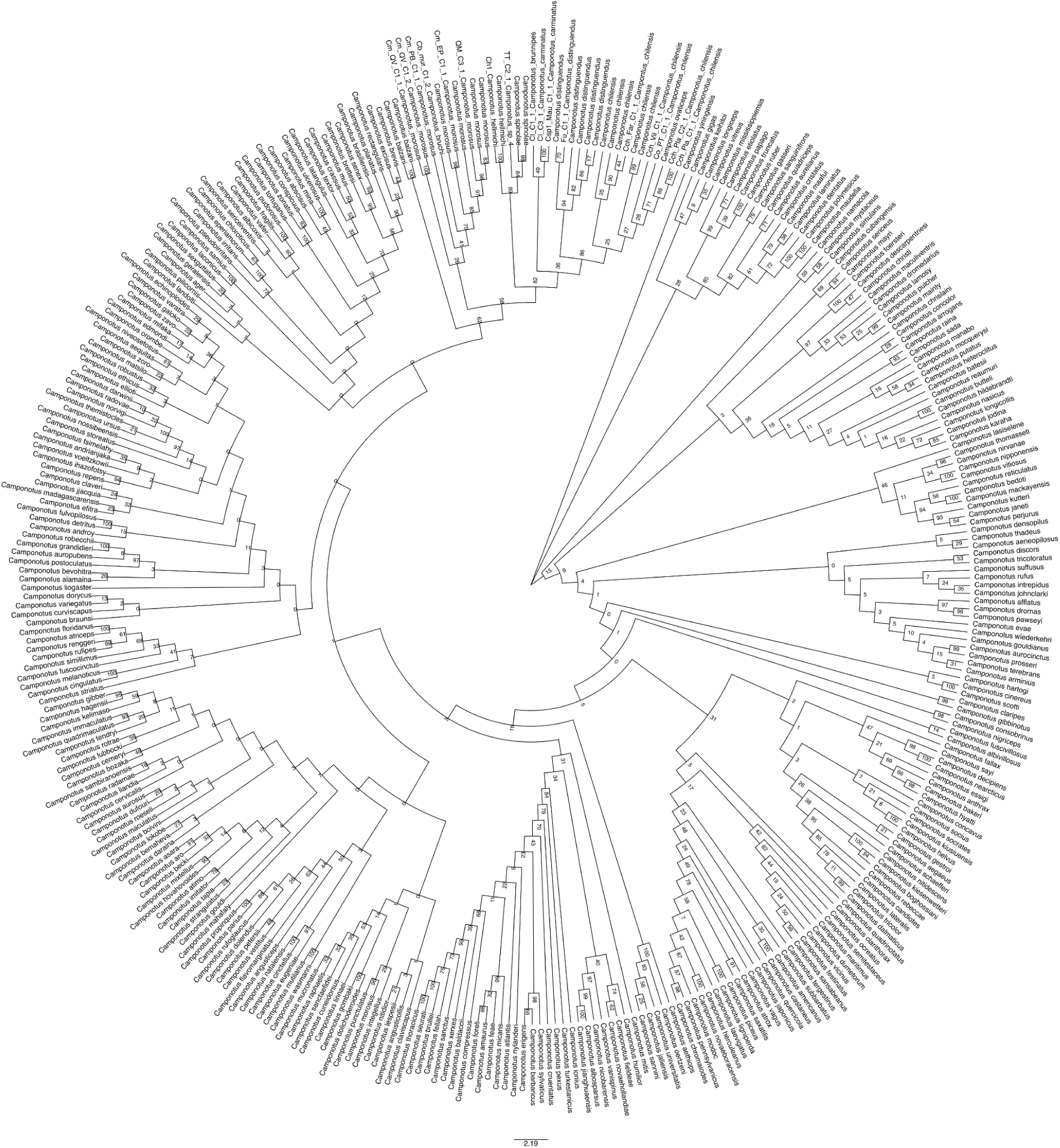
Global phylogeny of *Camponotus* species estimated from COI gene sequences representing 328. *Camponotus* species. Bootstrap values are shown numerically at each node.

**Figure S2:**
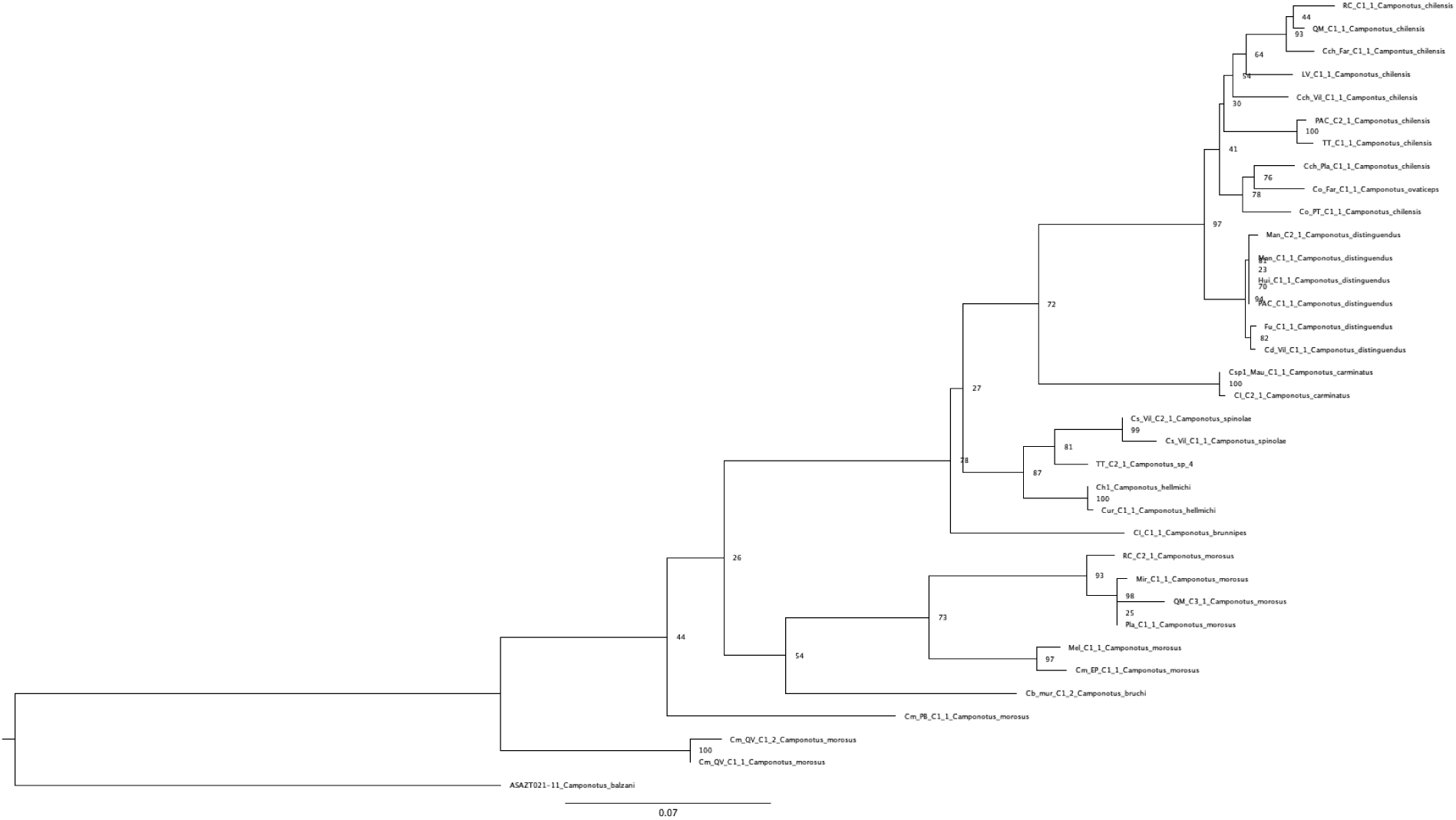
**Local phylogeny of *Camponotus* species present in Chile**, estimated from COI gene sequences. A total of 35 sequences were used for the estimation, including *Camponotus balzani*, which was used as the outgroup. Bootstrap values are shown numerically at each node.

**Figure S3:**
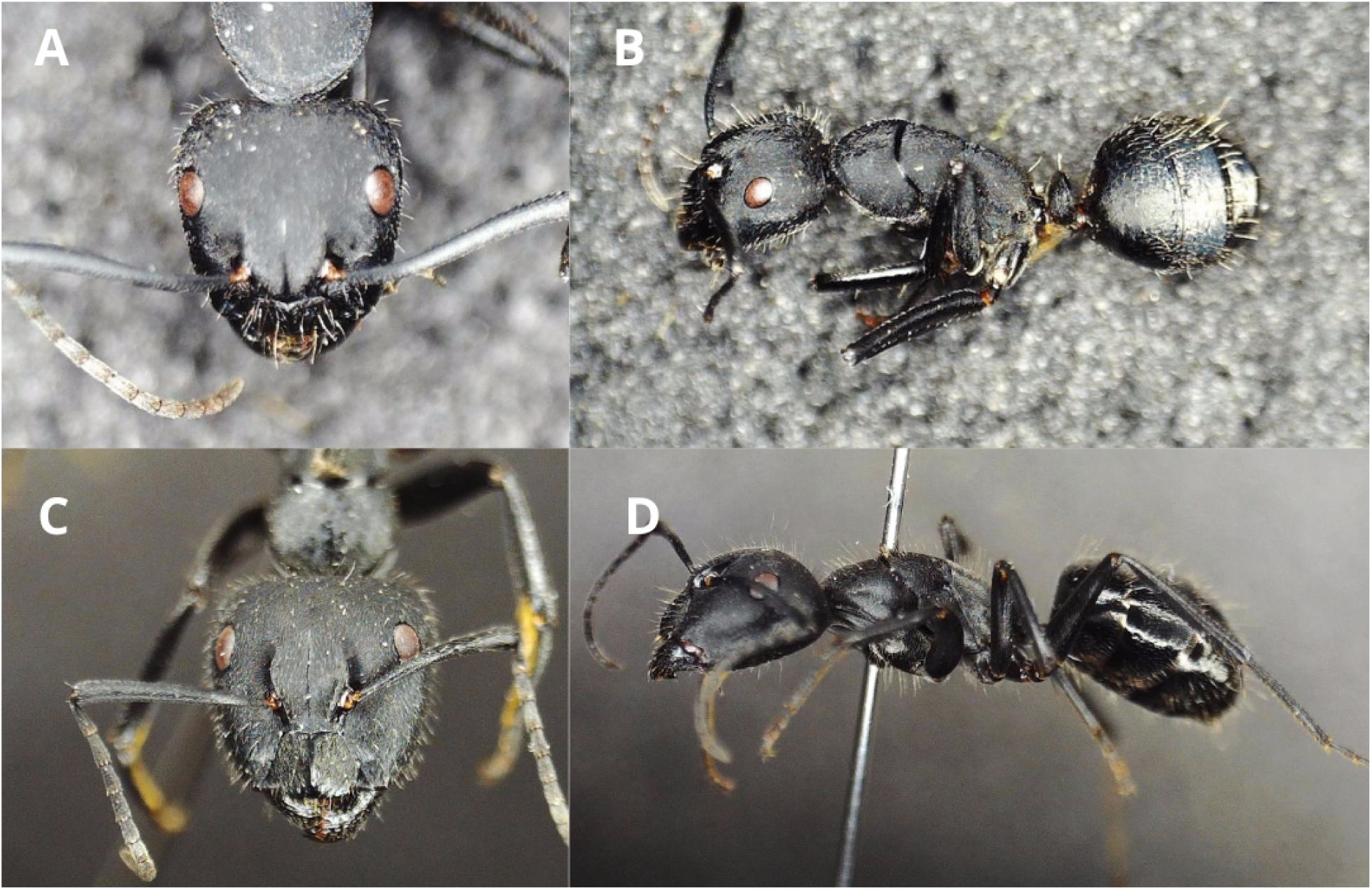
Photograph of TT_C2_1 (a and b) and *Camponotus distinguendus* (c and d). Cephalic view (a and c) and lateral view (b and d).

**Figure S4:**
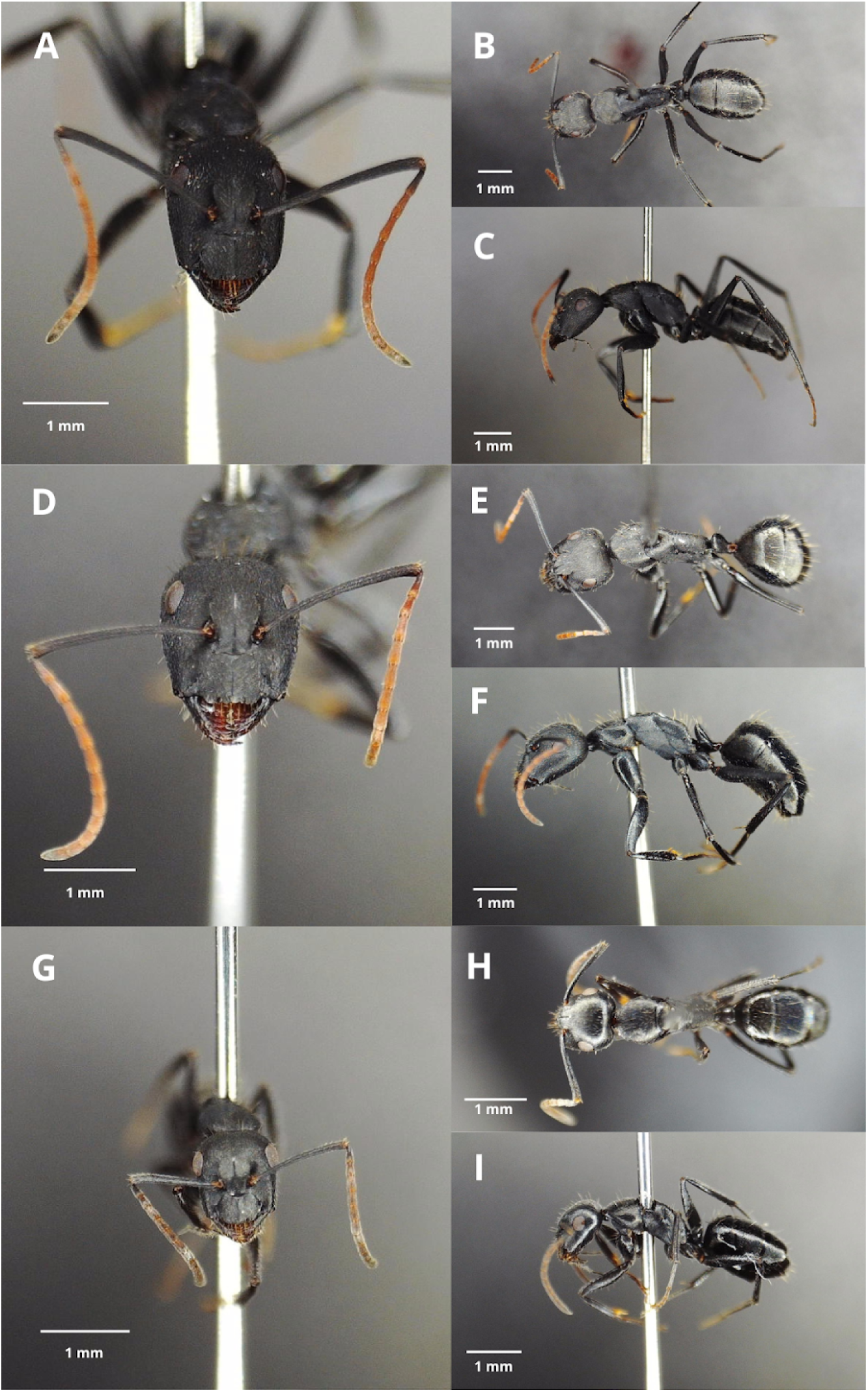
Photographs of *C. morosus* from different localities that exhibit genetic differences greater than 7%. (A, B, and C) Individual from the Andes Mountains of Santiago, locality QM_C3_1; (D, E, and F) Individual from Melipilla, locality Mel_C1_1; (G, H, and I) Individual from Laguna Verde, locality Cm_QV_C1_1. According to the key of Snelling and Hunt, all are categorized as *Camponotus morosus*.

**Figure S5:**
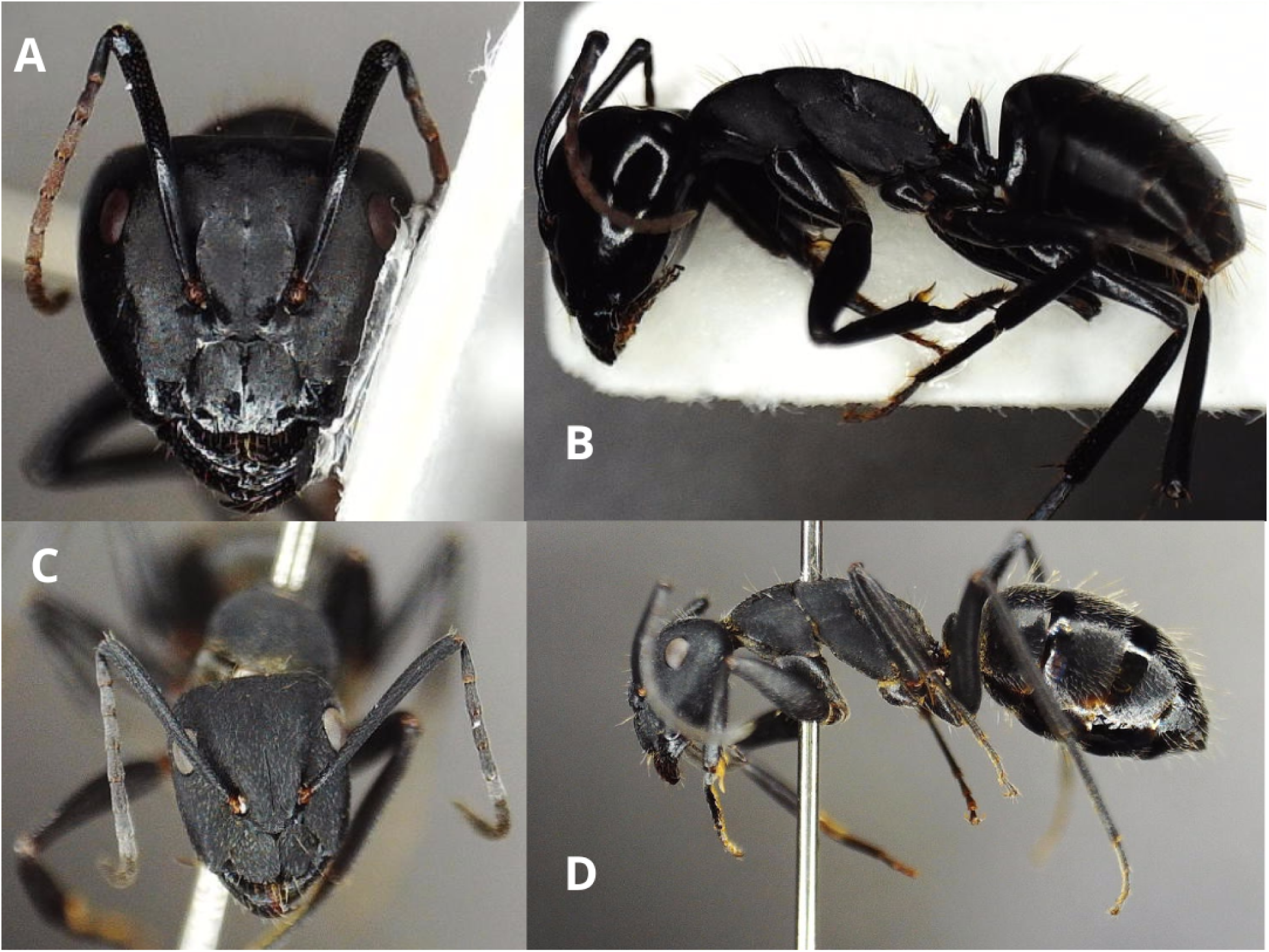
Photograph of *Camponotus* sp. 3 (a and b) and *Camponotus hellmichi* (c and d). Cephalic view (a and c) and lateral view (b and d).

